# Aerolysin enables modular, non-genetic functionalization of living cell surfaces

**DOI:** 10.64898/2026.08.28.746739

**Authors:** Andrew C.D. Lemmex, Mathew R. Pawlak, Wendy R. Gordon

## Abstract

Methods for installing synthetic functions on living cell surfaces provide powerful approaches for imaging, sensing, and manipulating cell behavior, but many require genetic modification of the target cell or chemical modification of the plasma membrane. Here, we repurpose the glycosylphosphatidylinositol-anchored protein (GPI-AP)-binding toxin aerolysin as a modular chassis for non-genetic cell-surface functionalization. We show that a non-cytotoxic, monomeric aerolysin mutant retains high-affinity and GPI-AP-dependent cell binding when genetically fused to diverse protein cargos. Fluorescent protein-aerolysin fusions robustly label multiple cell types and remain predominantly associated with the cell surface for at least 24 h, in contrast to wheat germ agglutinin, which is extensively internalized. Aerolysin can also be equipped with SpyTag/SpyCatcher to enable simple and modular assembly with independently expressed protein cargos. Importantly, aerolysin supports functional rather than solely optical modification of the cell surface: fusion to the proximity-labeling enzyme APEX2 enables cell surface protein biotinylation, while fusion to HUH endonuclease tags enables covalent attachment of synthetic DNA to living cells. Using this latter architecture, we developed a DNA hairpin sensor that converts cell-surface nuclease activity into a fluorescent signal and distinguishes cells with different levels of extracellular nuclease activity. Together, these results establish a non-cytotoxic and monomeric aerolysin mutant as a soluble adapter for installing proteins, enzymes, and programmable nucleic acids onto living cells without modification of the target-cell genome.

## INTRODUCTION

The plasma membrane provides a functional and dynamic interface through which cells sense, communicate with, and respond to their environment. The ability to experimentally modify this interface has therefore become an important strategy for interrogating and engineering cellular behavior. Genetically encoded receptors and surface tags provide precise means of introducing new functions onto cells, but require manipulation of the target-cell genome and are not readily applicable to all cell types or experimental settings. A growing set of non-genetic strategies instead modifies existing membrane components through direct chemical conjugation, metabolic glycan engineering, enzymatic remodeling, liposome fusion, or hydrophobic insertion of lipid-linked cargos. (19,20) Although powerful, these approaches often require chemical or metabolic preconditioning of the target cell, modify broad classes of membrane components, or depend on membrane-insertion and trafficking behavior. A simple soluble reagent capable of binding a broadly distributed endogenous cell-surface feature could provide a complementary strategy wherin functional proteins or synthetic molecules are prepared independently and subsequently added to cells to confer new surface properties.

Glycosylphosphatidylinositol-anchored proteins (GPI-APs) provide an attractive endogenous target for such an approach. GPI-APs comprise a diverse family of extracellular proteins tethered to the outer leaflet of the plasma membrane through a conserved glycolipid anchor and are broadly expressed on eukaryotic cells. Despite their diverse protein sequences and functions, all GPI-APs share features within the GPI anchor that are recognized by several microbial toxins. (1,2)

Aerolysin, a β-pore-forming toxin produced by *Aeromonas hydrophila*, binds GPI-APs with high affinity. In its native context, cell-surface binding is followed by oligomerization and formation of a membrane-spanning pore that compromises the cell membrane, resulting in cytotoxicity. However, structurally characterized aerolysin mutants separate GPI-AP recognition from pore formation. In particular, the T253/A300C aerolysin mutant retains cell binding while remaining monomeric and non-pore-forming. Chemically conjugated fluorescent derivatives of non-cytotoxic aerolysin have consequently been used to detect GPI-AP-deficient blood cells in paroxysmal nocturnal hemoglobinuria. We reasoned that this same property could be exploited more broadly: rather than using aerolysin solely to detect GPI-APs, aerolysin could serve as an adapter through which diverse molecular cargos are installed onto the surface of otherwise unmodified cells. (3–5) Here, we establish non-cytotoxic monomeric aerolysin as a modular chassis for cell-surface functionalization. We first demonstrate that aerolysin tolerates direct fusion to fluorescent and bioluminescent proteins without loss of GPI-AP-dependent cell binding and that a SpyTag/SpyCatcher architecture permits independently expressed cargos to be assembled with aerolysin in a plug-and-play manner. Aerolysin fusions label diverse cell types and, importantly, exhibit prolonged surface retention compared with wheat germ agglutinin. We then extend the platform beyond imaging by installing the enzymatically active proximity biotinylator APEX2 onto cell surfaces. Finally, by combining aerolysin with sequence-specific HUH endonuclease tags, we covalently install synthetic DNA onto living cells and use this architecture to construct a cell-surface nuclease biosensor. These results establish aerolysin as an easy to use and broadly applicable tool to target endogenous cell-surface glycoproteins to enable the use of and synthetic protein or nucleic-acid functionality.

## RESULTS

### Non-cytotoxic aerolysin functions as a high-affinity cell-surface anchor for protein cargos

To determine whether aerolysin would maintain its cell-surface anchoring properties while carrying an independently functional protein, we fused the fluorescent protein mNeonGreen to the N terminus of the monomeric T253/A300C aerolysin mutant (Fig. 1A). The resulting mNeonGreen-aerolysin fusion was expressed and purified from *E. coli* and compared with commercially available Alexa Fluor 488-conjugated aerolysin and Alexa Fluor 488-conjugated wheat germ agglutinin (WGA), a commonly used lectin that recognizes cell-surface glycans independently of the GPI anchor. (8)

**Figure 1.**
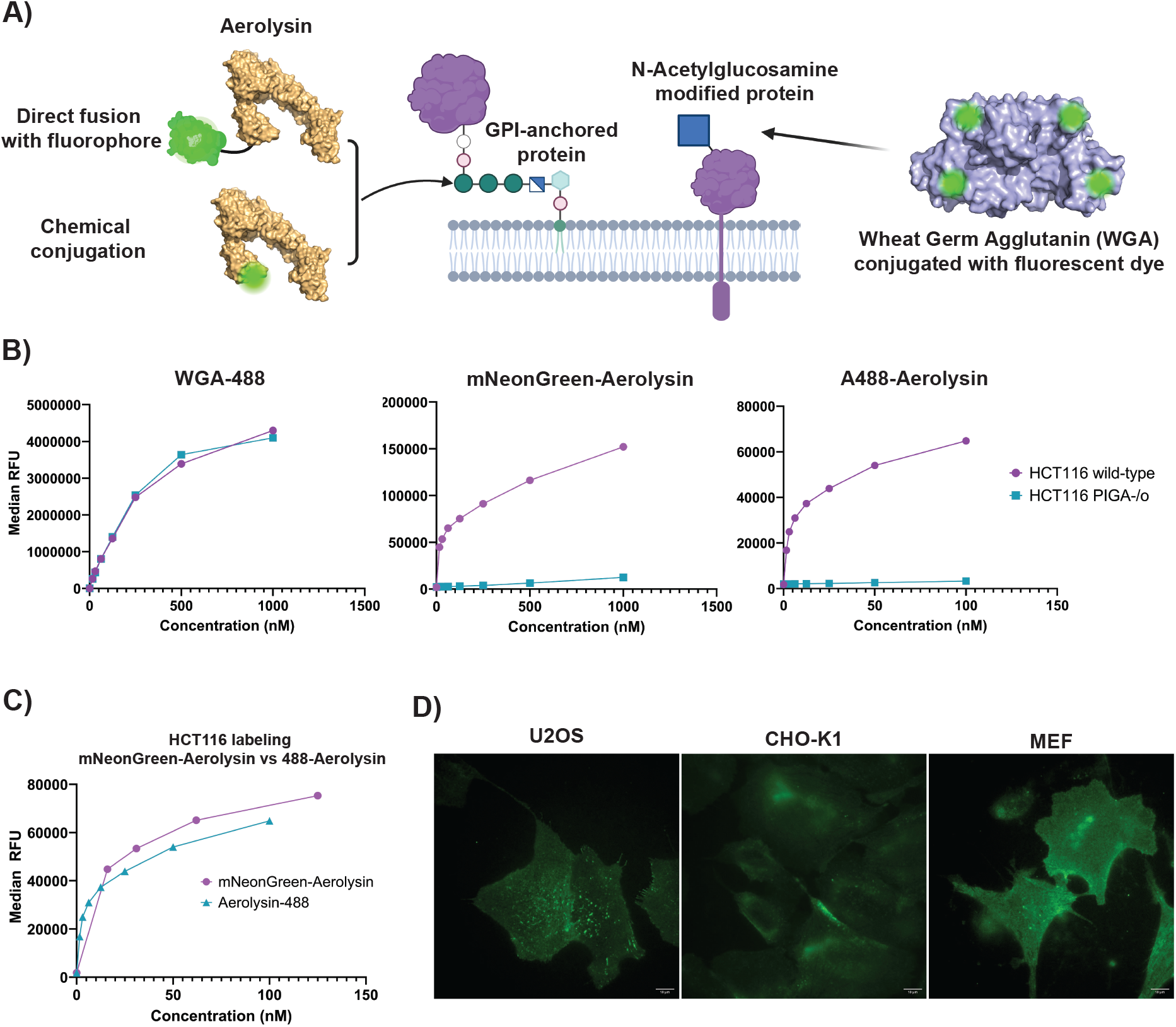
Aerolysin enables broad, GPI-anchor-dependent labeling of living cell surfaces. (A) Schematic comparing aerolysin-based cell-surface labeling with wheat germ agglutinin (WGA). Aerolysin can be genetically fused to a fluorescent protein or chemically conjugated to a fluorophore and targets glycosylphosphatidylinositol (GPI)-anchored proteins, whereas fluorescently conjugated WGA recognizes N-acetylglucosamine-containing glycoconjugates at the cell surface. (B) Concentration-dependent labeling of wild-type and PIGA-knockout HCT116 cells with WGA-488, mNeonGreen-aerolysin, or Alexa Fluor 488-conjugated aerolysin. WGA labeled both cell populations, whereas labeling by either aerolysin reagent was strongly reduced in PIGA-knockout cells, demonstrating dependence on GPI-anchored proteins. Median fluorescence intensity measured by flow cytometry is plotted as a function of labeling reagent concentration. (C) Comparison of concentration-dependent labeling of HCT116 cells with genetically encoded mNeonGreen-aerolysin and chemically labeled aerolysin-488. (D) Representative fluorescence microscopy images demonstrating mNeonGreen-aerolysin labeling of mouse embryonic fibroblasts (MEFs), CHO-K1, and U2OS cells. Together, these results establish aerolysin as a genetically and chemically adaptable cell-surface anchor that recognizes an endogenous, broadly distributed class of membrane proteins without requiring genetic modification of the target cell.

We first tested labeling of wild-type HCT116 cells and an isogenic PIGA-deficient HCT116 line (Fig. 1B-C). Since PIGA is required for an early step in GPI-anchor biosynthesis, its absence results in GPI-AP deficient cells. This provides a genetic control for GPI-AP-dependent binding. WGA robustly labeled both wild-type and PIGA-deficient cells. In contrast, both mNeonGreen-aerolysin and chemically labeled aerolysin strongly labeled wild-type HCT116 cells but showed little labeling of PIGA-deficient cells (Fig. 1B). Thus, fusion of a fluorescent protein to aerolysin preserved its dependence on GPI-AP expression. (9)

Titration of mNeonGreen-aerolysin revealed concentration-dependent labeling of wild-type HCT116 cells, with an apparent binding constant of approximately 110 nM. This was comparable to the binding behavior of commercially labeled aerolysin and confirmed that addition of a genetically encoded protein cargo did not substantially compromise cell-surface recognition (Fig. 1C). (10)

We next asked whether aerolysin labeling extended beyond HCT116 cells. mNeonGreen-aerolysin robustly labeled U2OS, CHO-K1, and mouse embryonic fibroblast cells, although the spatial distribution of fluorescence differed between cell types (Fig. 1D). Together, these results indicate that non-cytotoxic aerolysin can function as a genetically encoded cell-surface anchoring domain across diverse cellular backgrounds.

### A modular SpyTag/SpyCatcher architecture enables plug-and-play aerolysin functionalization

Direct protein fusion provides a simple route for producing aerolysin-cargo proteins but requires each cargo to tolerate expression and purification in the same context as aerolysin. We therefore sought to develop a modular architecture in which aerolysin and its cargo could be produced independently and subsequently assembled.

We evaluated both split-intein-mediated protein ligation and the SpyTag/SpyCatcher system (Fig. 2A and Fig. S1A). Both approaches generated aerolysin-cargo conjugates in vitro. However, SpyTag/SpyCatcher assembly was particularly robust and proceeded without the reducing conditions required for the split-intein reaction. Therefore the SpyTag/Spycatcher system was selected for further characterization. (11–13)

**Figure 2.**
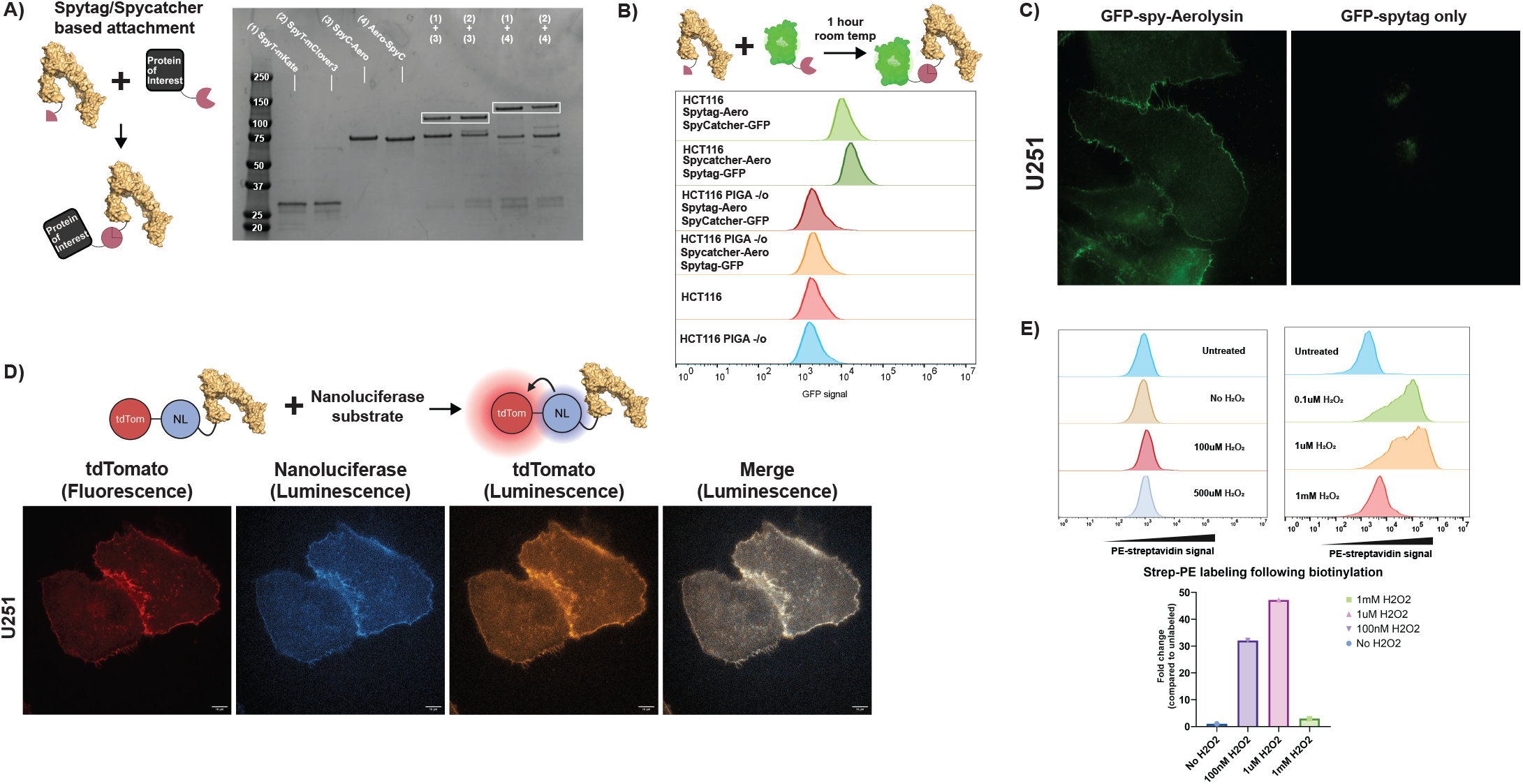
Aerolysin supports modular installation of diverse functional protein cargos on living cell surfaces. **(A)** Schematic of the SpyTag/SpyCatcher strategy for modular assembly of independently expressed proteins of interest with aerolysin. **(B)** SDS-PAGE analysis demonstrating covalent assembly of SpyTag/SpyCatcher-aerolysin constructs with protein cargo. Boxed bands indicate products corresponding to the assembled complexes. **(C)** SpyTag/SpyCatcher-mediated assembly of GFP with aerolysin enables GPI-anchor-dependent cell labeling. HCT116 wild-type and PIGA-knockout cells were incubated with the indicated GFP/SpyTag/SpyCatcher constructs and analyzed by flow cytometry. **(D)** Representative fluorescence microscopy of cells labeled with GFP assembled with aerolysin through SpyTag/SpyCatcher compared with GFP-SpyTag alone. **(E)** Aerolysin fused to the Nano-lantern bioluminescent reporter retains both cell-surface targeting and reporter activity. Schematic illustrates substrate-dependent NanoLuc luminescence and BRET to tdTomato; representative images show tdTomato fluorescence, NanoLuc luminescence, tdTomato acceptor luminescence, and merged signal in U251 cells labeled with the fusion protein. **(F)** APEX2-aerolysin enables enzymatic labeling at the cell surface. Cells labeled with APEX2-aerolysin were subjected to proximity biotinylation under the indicated H_2_O_2_ conditions, and biotinylation was quantified by PE-streptavidin flow cytometry. Representative fluorescence distributions and quantification relative to untreated cells are shown. Together, these results demonstrate that aerolysin can accommodate both modularly assembled and directly fused cargos spanning fluorescent, bioluminescent, and enzymatic functions while retaining cell-surface targeting and cargo activity.

SpyCatcher could be positioned at either terminus of aerolysin and efficiently reacted with SpyTag-containing protein partners, including multiple fluorescent proteins (Fig. 2B). To determine whether this additional protein architecture altered cell binding, we assembled GFP-SpyTag with SpyCatcher-aerolysin and incubated the resulting complexes with HCT116 cells. The assembled GFP-aerolysin conjugates robustly labeled wild-type cells by flow cytometry while retaining low labeling of PIGA-deficient cells (Fig. 2C). Fluorescence microscopy similarly revealed strong plasma-membrane-associated GFP signal with little background from GFP-SpyTag alone.These results demonstrated that aerolysin retained its GPI-AP binding function without requiring direct genetic fusion to its functional cargo. Instead, a common aerolysin adapter can be assembled with independently produced proteins, providing a modular route for expanding the range of molecules that can be recruited to the cell surface.

### Aerolysin compatible with cell-surface bioluminescence

Aerolysin was also compatible with imaging modalities beyond conventional fluorescence. Fusion of aerolysin to a tdTomato-NanoLuc Nano-lantern generated robust fluorescence and bioluminescence from labeled U251 cells following addition of luciferase substrate (Fig. 2D). Thus, the aerolysin anchoring domain can support diverse optical reporters while retaining cell-surface targeting. (14)

### Aerolysin can install enzymatic activity on the extracellular cell surface

We next asked whether aerolysin could deliver a functional enzyme rather than an optical reporter. We selected APEX2, an engineered peroxidase widely used for proximity-dependent biotinylation, and fused it to the N terminus of aerolysin. Following incubation of U2OS cells with APEX2-aerolysin, cells were exposed to biotin-phenol and hydrogen peroxide and biotinylation was detected using fluorescent streptavidin. (15)

APEX2-aerolysin produced readily detectable cell-associated biotinylation, demonstrating that both the aerolysin targeting domain and APEX2 enzymatic domain remained functional in the fusion protein (Fig. 2E). Interestingly, labeling efficiency was strongly dependent on hydrogen peroxide concentration. Hydrogen peroxide concentrations greater than approximately 1 μM reduced labeling under the conditions used here with the effect increasing with greater concentraitons of hydrogen peroxide, whereas 1 μM and 100 nM hydrogen peroxide generated substantially stronger streptavidin signals (Fig. 2E).

These results demonstrate that aerolysin can install catalytically active protein machinery onto the extracellular surface of living cells. Thus, the platform is not limited to passive labeling but can confer new biochemical activity on the target-cell surface.

### Aerolysin enables versatile and persistent labeling of living cells

We next investigated properties of aerolysin that could be useful for live-cell labeling. Fluorescent aerolysin constructs labeled multiple cell types and could accommodate fluorescent proteins with distinct spectral properties. Interestingly, aerolysin labeling also revealed cell-type-dependent distributions of GPI-AP-containing membrane domains. For example, mNeonGreen-aerolysin produced relatively diffuse surface staining in mouse embryonic fibroblasts but more punctate patterns in U2OS cells (Fig. 1D).

To directly compare aerolysin with an established reagent for labeling the extracellular cell surface, we colabeled cells with mScarlet-aerolysin and fluorescent WGA. The two reagents produced partially overlapping but clearly distinct distributions. In BT-474 cells, WGA displayed prominent punctate staining whereas aerolysin labeling was more diffuse. IPN-Schwann cells showed a different pattern, including pronounced aerolysin signal at the cell periphery (Fig. 3B), consistent with the two reagents recognizing distinct classes of cell-surface glycoconjugates.

**Figure 3.**
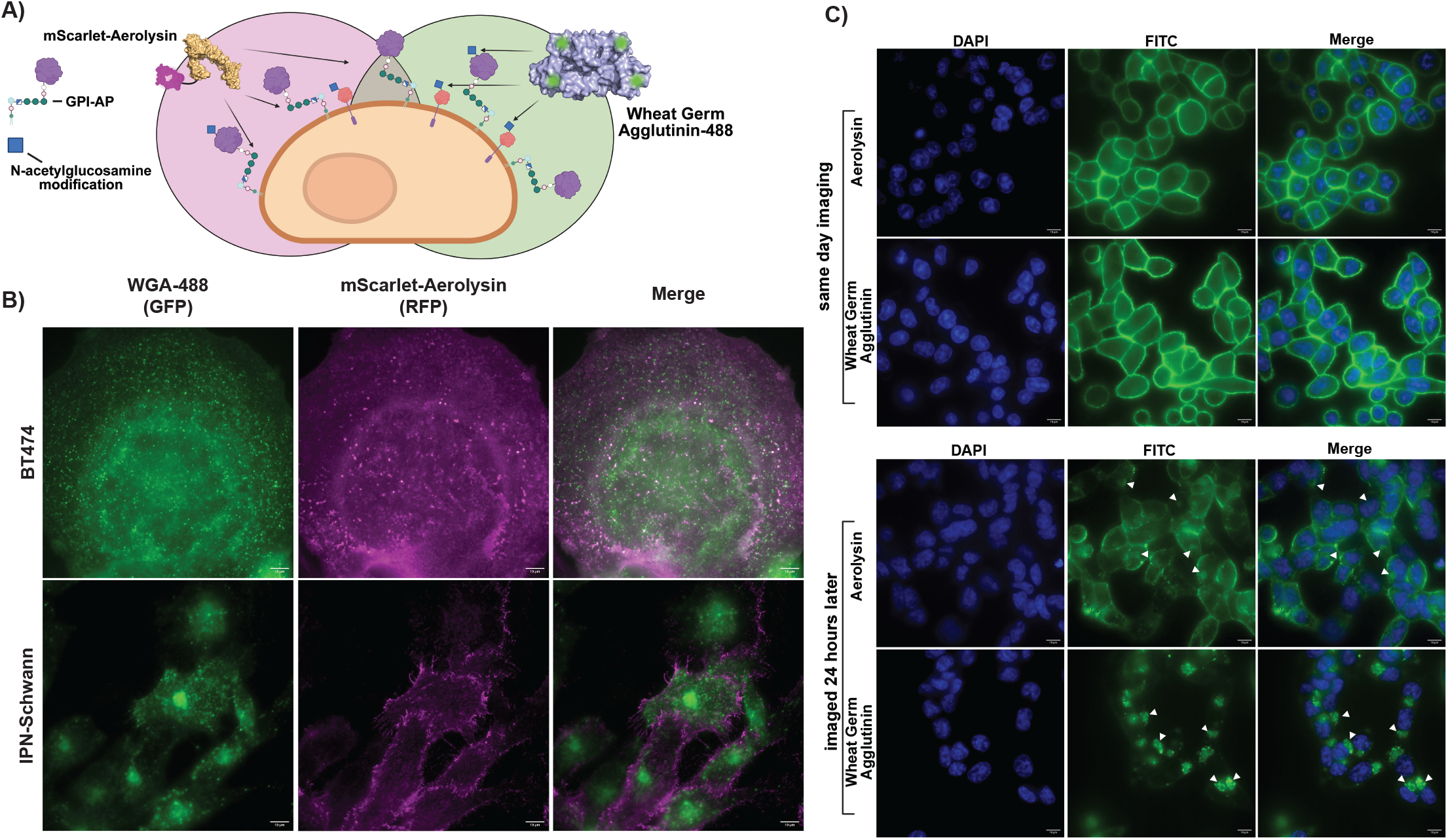
Aerolysin provides spatially distinct and persistent labeling of the cell surface. **(A)** Schematic illustrating the distinct cell-surface targets of aerolysin and wheat germ agglutinin (WGA). mScarlet-aerolysin binds glycosylphosphatidylinositol-anchored proteins (GPI-APs), whereas fluorescent WGA recognizes N-acetylglucosamine-containing glycoconjugates, enabling simultaneous visualization of two distinct classes of cell-surface components. **(B)** Representative fluorescence microscopy of BT474 and IPN-Schwann cells co-labeled with WGA-488 (green) and mScarlet-aerolysin (magenta). Aerolysin and WGA exhibit distinct spatial distributions at the plasma membrane, with aerolysin displaying prominent punctate and membrane-associated labeling that only partially overlaps with WGA. **(C)** Persistence of aerolysin- and WGA-mediated labeling over time. Cells were labeled with fluorescent aerolysin or WGA and imaged either on the day of labeling or 24 h later. Both reagents show prominent cell-surface labeling immediately following treatment. Aerolysin-associated fluorescence remains detectable 24 h after labeling, including in discrete punctate structures (arrowheads), demonstrating persistence of the aerolysin-delivered fluorescent signal over time. Nuclei were stained with DAPI. Scale bars are as indicated.

We next compared the persistence of the two labels. Immediately after a brief labeling and wash step, both WGA and mNeonGreen-aerolysin predominantly labeled the HCT116 cell surface. Twenty-four hours later, aerolysin fluorescence remained readily detectable at the cell surface, with comparatively limited intracellular signal. In contrast, surface-associated WGA fluorescence was markedly reduced and the remaining signal appeared predominantly intracellular (Fig. 3C). These observations indicate that aerolysin can provide prolonged labeling of the extracellular membrane compared with WGA under these conditions.

### HUH-aerolysin covalently installs synthetic DNA onto living cells

The ability to recruit synthetic nucleic acids to cells would substantially expand the chemical functionality accessible through the aerolysin platform. We therefore fused aerolysin to an HUH endonuclease tag. HUH endonucleases recognize defined single-stranded DNA sequences and form a covalent phosphotyrosine linkage with their DNA substrate, enabling direct and sequence-dependent protein-DNA conjugation (Fig. 4A). (16)

**Figure 4.**
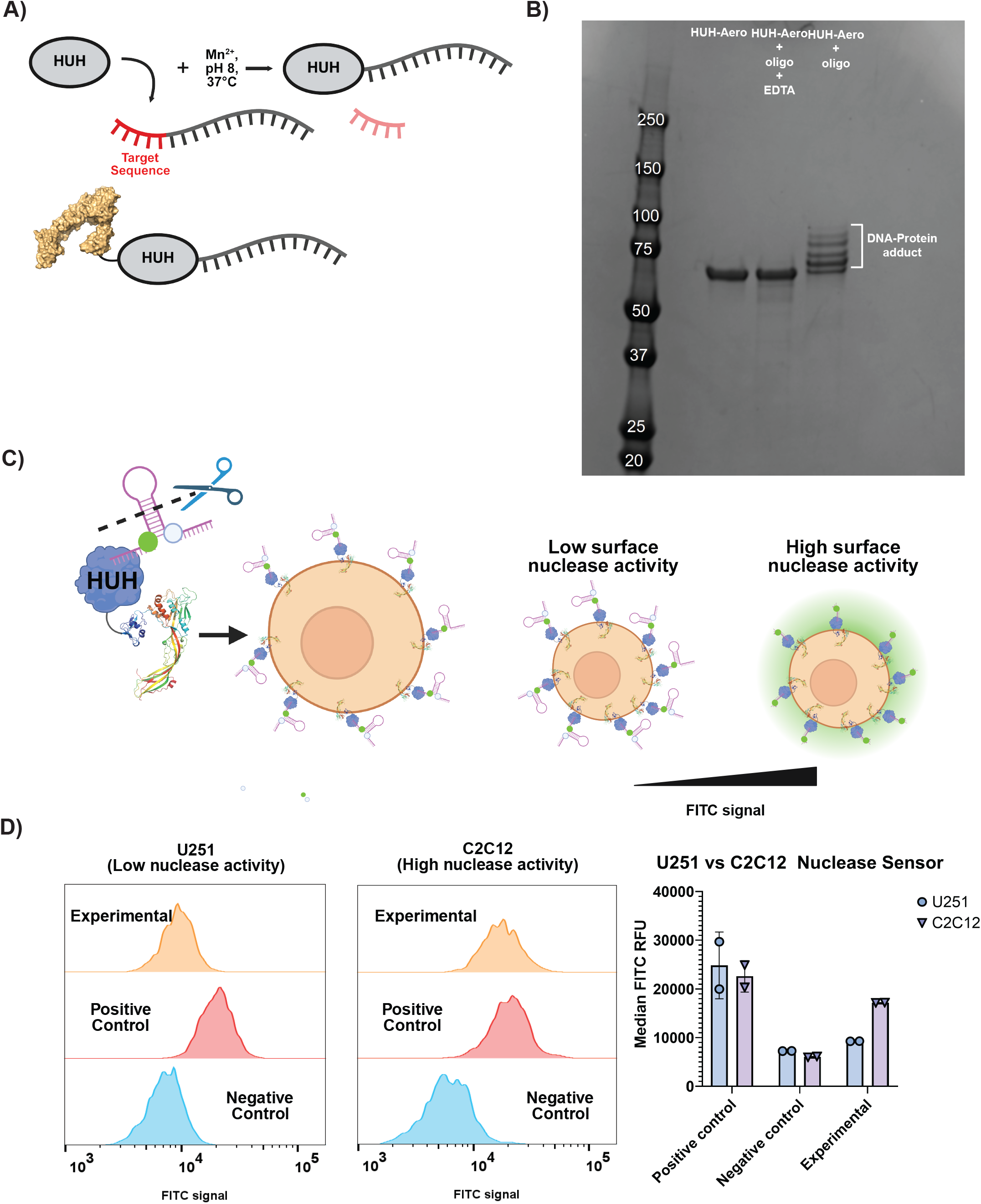
HUH-aerolysin enables programmable DNA display and cell-surface nuclease sensing. **(A)** Schematic of HUH-mediated covalent protein–DNA conjugation. HUH endonucleases recognize a specific single-stranded DNA sequence and, in the presence of Mn^2+^, form a covalent phosphotyrosine linkage with the DNA. Fusion of the HUH domain to aerolysin enables covalently attached DNA cargos to be displayed on living cells through aerolysin binding to GPI-anchored proteins. **(B)** SDS-PAGE analysis of HUH-aerolysin conjugation to its cognate oligonucleotide. Formation of the covalent DNA–protein adduct produces the indicated mobility shift; EDTA inhibits the metal-dependent conjugation reaction. **(C)** Design of the RASUN (aerolysin surface nuclease) sensor. A fluorophore-containing DNA nuclease sensor is covalently attached to HUH-aerolysin and displayed on the cell surface. Cleavage of the DNA sensor by extracellular nucleases separates the fluorophore from its quenched state, producing a fluorescence signal proportional to cell-surface nuclease activity. **(D)** Flow-cytometric detection of cell-surface nuclease activity in U251 and C2C12 cells. Representative FITC fluorescence distributions are shown for experimental nuclease sensors together with positive and negative controls. Quantification of median FITC fluorescence demonstrates greater sensor activation in C2C12 cells than in U251 cells, consistent with higher cell-surface nuclease activity. Points represent independent experiments; bars show [median ± SEM].

An aerolysin fusion containing a porcine circovirus-derived HUH tag readily formed covalent DNA-protein adducts under standard HUH reaction conditions. Multiple products were observed with a DNA substrate containing multiple compatible recognition sites, whereas chelation of the required divalent metal with EDTA inhibited conjugate formation (Fig. 4B). These results demonstrated that both aerolysin and the HUH domain tolerate fusion without eliminating HUH-mediated DNA conjugation.

### Aerolysin-anchored DNA enables sensing of cell-surface nuclease activity

We next used HUH-aerolysin to determine whether cell-anchored DNA could function as a biosensor rather than simply as a label. We designed a DNA hairpin containing a fluorophore-quencher pair such that the intact hairpin exhibited low fluorescence. The sensor was covalently linked to HUH-aerolysin near the fluorophore-containing end of the DNA. We reasoned that cleavage of the surface-displayed DNA by extracellular nucleases would separate the fluorophore from its quencher and increase fluorescence (Fig. 4C). We termed this approach RApid Sensing of NUcleases (RASUN).

As an initial test, we compared U251 and C2C12 cells, which exhibited relatively low and high cell-surface nuclease activity, respectively. Cells were labeled with HUH-aerolysin carrying the quenched DNA hairpin and incubated for 90 min before analysis by flow cytometry. The sensor produced different fluorescence responses in the two cell types, with C2C12 cells generating substantially greater signal than U251 cells (Fig. D). Thus, aerolysin-anchored DNA can convert differences in extracellular enzymatic activity into a cell-associated fluorescent readout.

Together, these experiments establish a progression from cell-surface targeting to cell-surface programming: aerolysin recruits the HUH-DNA conjugate to endogenous GPI-APs, while the sequence and structure of the attached DNA determine the biochemical information that can be read from the cell.

## DISCUSSION

Here, we repurpose a non-cytotoxic monomeric mutant of the bacterial toxin aerolysin as a modular platform for installing synthetic functionality onto living cell surfaces. The central feature of this strategy is that the target cell itself does not need to be genetically engineered to encode a receptor, tag, or synthetic binding site.

Instead, aerolysin exploits broadly distributed endogenous GPI-APs as docking sites for soluble protein reagents produced independently of the cell. By changing the molecular cargo associated with aerolysin, the same targeting mechanism can support fluorescent and bioluminescent imaging, extracellular enzymatic activity, or display of programmable DNA.

Several features make aerolysin attractive as a cell-surface anchoring chassis. First, fusion of mNeonGreen to the monomeric T253/A300C mutant preserved GPI-AP-dependent binding: labeling was robust in wild-type HCT116 cells but strongly reduced in PIGA-deficient cells. The fusion protein also exhibited concentration-dependent binding comparable to chemically labeled aerolysin and labeled multiple unrelated cell types. These observations suggest that the aerolysin scaffold tolerates substantial N-terminal modification without disrupting recognition of its endogenous cell-surface targets.

Second, aerolysin provides flexibility in how functional cargos are incorporated. Direct genetic fusion is straightforward for proteins that can be co-expressed with aerolysin, whereas the SpyTag/SpyCatcher architecture allows aerolysin and its cargo to be produced independently and assembled after purification. The latter feature may be particularly useful for cargos requiring different expression systems or purification conditions. More broadly, this modularity allows a common aerolysin reagent to serve as an adapter rather than requiring the complete cell-targeting protein to be redesigned for every application, substantially reducing the burden for reagent preparation.

Aerolysin also displayed properties useful for longitudinal cell labeling. In our experiments, fluorescent aerolysin remained predominantly associated with the cell surface 24 h after labeling, whereas WGA became extensively internalized over the same period. Because aerolysin and WGA recognize distinct classes of glycoconjugates, this difference likely reflects their respective membrane targets and trafficking pathways rather than a universal advantage of one reagent over the other. Nevertheless, the persistence of aerolysin labeling suggests that it may be particularly useful for experiments requiring sustained access to the extracellular membrane.

Importantly, the aerolysin platform is not limited to optical labeling. APEX2 retained enzymatic activity when fused to aerolysin and enabled biotinylation at the cell surface. This result demonstrates that aerolysin can position a catalytically active protein at the plasma membrane and suggests potential applications in extracellular proximity labeling or interrogation of GPI-AP-associated membrane environments. Additional optimization and proteomic characterization will be required to establish the spatial selectivity and proteome coverage of APEX2-aerolysin, and the present experiments should therefore be viewed primarily as demonstration of enzymatic cargo delivery rather than a fully developed surface-proteomics method. (17)

The HUH-aerolysin architecture extends this concept by separating cell targeting from molecular programming. HUH tags form covalent bonds with sequence-defined single-stranded DNA substrates, allowing a single recombinant HUH-aerolysin protein to be conjugated to different synthetic oligonucleotides. The resulting DNA-protein conjugates efficiently label cells, providing a simple route for decorating an unmodified plasma membrane with synthetic nucleic acids. Because oligonucleotides can encode recognition elements, fluorophores, quenchers, barcodes, molecular switches, and other functional structures, this architecture potentially provides substantially greater programmability than a fixed protein fusion.

RASUN illustrates one application of this programmability. By attaching a quenched DNA hairpin to HUH-aerolysin, extracellular nuclease activity is converted into a fluorescent signal associated with the labeled cell. The sensor distinguished U251 and C2C12 cells with different nuclease activities and provides a foundation for determining whether cell-surface nuclease activity changes dynamically with cell state. (18)

The aerolysin platform also has limitations. Because labeling depends on endogenous GPI-AP abundance and composition, labeling density is expected to vary among cell types and cell states. Aerolysin does not provide molecular specificity for an individual GPI-AP and should instead be viewed as targeting a broadly distributed class of endogenous membrane anchors. In addition, despite the use of non-pore-forming aerolysin mutants, the consequences of prolonged or high-density occupancy of GPI-APs should be evaluated for applications in which preservation of native membrane organization or signaling is critical. These characteristics distinguish aerolysin from genetically encoded systems that provide precise control over the identity and abundance of an engineered surface receptor.

Rather than replacing such approaches, aerolysin provides a complementary strategy for situations in which rapid and non-genetic modification is advantageous. Functional aerolysin reagents can be produced in advance, added directly to cells, and removed by washing, making the approach particularly attractive for primary cells, transient experiments, or comparisons across multiple cell types where genetic manipulation would be cumbersome. (19,20)

More broadly, this work suggests that the natural targeting capabilities of non-cytotoxic toxin derivatives can be repurposed as tools for cell engineering. Toxins have evolved high-affinity recognition of abundant extracellular features in order to deliver biological activity to cells. Separating target recognition from toxicity converts this evolutionary specialization into a potentially useful molecular interface. Aerolysin provides one example in which an endogenous membrane feature can be exploited to install synthetic proteins and nucleic acids onto living cells. Expanding this strategy to other non-cytotoxic toxin scaffolds with distinct cell-surface specificities could ultimately create a toolbox of soluble adapters for non-genetic programming of living cell surfaces.

## CONCLUSION

Non-cytotoxic aerolysin provides a simple molecular bridge between endogenous GPI-anchored proteins and exogenously produced functional cargos. By combining aerolysin with fluorescent proteins, modular protein-conjugation systems, active enzymes, and HUH-mediated DNA conjugation, we demonstrate that living cells can be labeled, functionalized, and equipped with synthetic biosensors without genetic modification of the target cell. The resulting platform transforms aerolysin from a GPI-AP detection reagent into a modular chassis for cell-surface engineering and provides a foundation for programmable interrogation of dynamic extracellular biology.

## EXPERIMENTAL METHODS

### Cell culture

HCT116 wild-type and PIGA-deficient HCT116 cells were cultured in McCoy’s 5A medium. U251, U2OS, IPN-Schwann, C2C12, and mouse embryonic fibroblast (MEF) cells were maintained in DMEM. CHO-K1 and BT-474 cells were maintained in F-12K medium. Media were supplemented with 10% fetal bovine serum and GlutaMAX where appropriate. Cells were maintained at 37°C in a humidified atmosphere containing 5% CO_2_. Unless otherwise indicated, aerolysin labeling and other cell treatments were performed in the corresponding complete culture medium.

## METHOD DETAILS

### Plasmid construction

Recombinant aerolysin constructs were cloned into the pTD68 bacterial expression vector containing an N-terminal 6×His-SUMO purification tag. DNA inserts were synthesized as gene blocks (Integrated DNA Technologies) and assembled into BamHI/XhoI-linearized pTD68 using In-Fusion HD cloning (Takara Bio) according to the manufacturer’s instructions. Assembly products were transformed into Stellar competent *E. coli* (Takara Bio), selected on LB agar containing 100 μg/mL ampicillin, and plasmids from individual colonies were verified by DNA sequencing. Sequence-verified constructs were subsequently transformed into BL21 *E. coli* for protein expression. SpyTag-mKate2 and SpyTag-mClover3 constructs in pET28 were obtained from Addgene (plasmids 133452 and 133453).

### Recombinant protein expression and purification

Aerolysin fusion proteins were expressed in BL21 *E. coli*. An overnight 10-mL LB culture containing ampicillin was used to inoculate 1 L LB containing ampicillin. Cultures were grown at 37°C to an OD_600_ of 0.8–1.0, cooled at 4°C for 1 h, and induced with 1 mM IPTG. Protein expression proceeded overnight at 18°C. Cells were harvested by centrifugation at 4,000 × *g* for 30 min at 4°C.

Cell pellets were resuspended in 30 mL lysis buffer (50 mM Tris, 500 mM NaCl, pH 8.0) supplemented with EDTA-free protease inhibitor (Pierce) and lysed by four 1-min rounds of sonication at 30% duty cycle. Lysates were clarified by centrifugation at 24,000 × *g* for 1 h, and imidazole was added to the supernatant to 20 mM. Proteins were captured using HisPur Ni-NTA resin (Thermo Fisher Scientific, 88222), washed with 50 mM Tris, 500 mM NaCl, 30 mM imidazole, pH 8.0, and eluted with 50 mM Tris, 500 mM NaCl, 300 mM imidazole, pH 8.0.

The N-terminal 6×His-SUMO tag was removed by addition of ULP1 during overnight dialysis against 50 mM Tris, 500 mM NaCl, 15 mM imidazole, pH 8.0. Cleaved protein was separated from His-tagged species by reverse Ni-NTA purification, and the flow-through containing the protein of interest was concentrated using a 30-kDa MWCO centrifugal concentrator. Proteins were further purified by size-exclusion chromatography on a Superdex 75 Increase 10/300 GL column equilibrated in 25 mM HEPES, 300 mM NaCl, pH 7.8. Fractions containing purified protein were identified by SDS-PAGE, pooled, concentrated, aliquoted, and stored at −80°C.

### SpyTag/SpyCatcher-mediated protein assembly

SpyTag/SpyCatcher conjugation reactions were performed in 25 mM HEPES, 200 mM NaCl, pH 7.4. Unless otherwise indicated, the non-aerolysin component was supplied at approximately 1.5-fold molar excess relative to the aerolysin-containing partner (typically 15 μM and 10 μM, respectively). Reactions were incubated for 1 h at room temperature and either used immediately or stored at −80°C. Formation of covalent conjugates was assessed by SDS-PAGE. For cell-labeling experiments, assembled products were treated as quantitatively reacted based on the near-complete conversion observed by SDS-PAGE.

For GFP labeling experiments, assembled GFP-SpyTag/SpyCatcher-aerolysin complexes were diluted to 200 nM and incubated with cells for 30 min before washing and analysis by fluorescence microscopy or flow cytometry. HCT116 wild-type and PIGA-deficient cells were used to assess retention of GPI-AP-dependent binding.

### Split-intein-mediated protein ligation

Split-intein reactions were performed using complementary CfaC- and CfaN-tagged proteins, generally at a 1:1 molar ratio, in 20 mM HEPES, 150 mM NaCl, pH 7.4. TCEP was included as a reducing agent; 2 mM TCEP was sufficient for routine reactions unless otherwise indicated. Reactions were typically incubated for 15 min at 30°C and analyzed by SDS-PAGE. For optimization experiments, TCEP concentration, reaction temperature, and incubation time were varied as indicated in Figure S2.

### Aerolysin-mediated cell labeling

Unless otherwise specified, aerolysin proteins were diluted to the desired concentration in complete cell-culture medium prewarmed to 37°C. Culture medium was removed and replaced with medium containing the labeling reagent, and cells were incubated for 30 min at 37°C. Cells were then washed repeatedly to remove unbound reagent. For live-cell imaging, cells were washed into FluoroBrite medium containing GlutaMAX before imaging.

For concentration-response experiments, wild-type and PIGA-deficient HCT116 cells were incubated with increasing concentrations of mNeonGreen-aerolysin, Alexa Fluor 488-conjugated aerolysin (Cedarlane, 4111015), or Alexa Fluor 488-conjugated wheat germ agglutinin (WGA; Biotium, 29022), followed by washing and flow-cytometric analysis. Median cellular fluorescence was plotted as a function of reagent concentration and fit using a nonlinear binding model to estimate apparent binding constants. The thesis reports an apparent binding constant of approximately 110 nM for mNeonGreen-aerolysin.

### Long-term cell-surface labeling

To compare the persistence of aerolysin and WGA labeling, HCT116 cells were incubated with 150 nM mNeonGreen-aerolysin or WGA-488 for 5 min and washed to remove unbound reagent. Cells were imaged immediately after labeling, returned to standard culture conditions, and imaged again 24 h later.

### Fluorescence and bioluminescence microscopy

Unless otherwise indicated, fluorescence microscopy was performed using an Olympus IX83 inverted microscope equipped with a motorized stage and stage-top incubator. Images were acquired using a 100× oil-immersion objective and processed using ImageJ.

For Nano-lantern imaging, cells were labeled and washed as described above. NanoLuc substrate (Promega, N2011) was prepared according to the manufacturer’s instructions and added immediately before bioluminescence imaging.

### Flow cytometry

Flow-cytometric measurements were performed using a BD Accuri C6 Plus flow cytometer. Following experimental treatment, adherent cells were washed two to three times with PBS and detached using a minimal volume of trypsin (Gibco, 25200056; approximately 30 μL per well for a 96-well plate). Cells were incubated at 37°C for 5 min and resuspended in flow buffer consisting of PBS containing 1% BSA and 1 mM EDTA. Cell suspensions were transferred to tubes and immediately analyzed by flow cytometry.

### APEX2-aerolysin proximity biotinylation

Cells were seeded one day before labeling at approximately one-third to one-half of the density expected at confluence. The following day, cells were incubated with 500 nM APEX2-aerolysin in complete culture medium for 1 h. Cells were washed with PBS and incubated with biotin-phenol in culture medium for 15 min. Biotinylation was initiated by addition of biotin-phenol and H_2_O_2_ in PBS. H_2_O_2_ concentrations were varied as indicated; concentrations of 1 μM or lower produced the strongest labeling under the conditions tested.

Reactions were quenched using PBS containing 5 mM Trolox and 10 mM sodium ascorbate, prewarmed to 37°C, followed by three washes with quench solution. Biotinylated proteins were detected by incubation with PE-conjugated streptavidin (BioLegend, 405204) diluted 1:500 in culture medium for 30 min at 37°C. Cells were washed three times with PBS and analyzed by fluorescence microscopy or flow cytometry.

### HUH-mediated protein–DNA conjugation

HUH-aerolysin constructs containing a porcine circovirus-derived HUH endonuclease were reacted with single-stranded DNA substrates containing the appropriate HUH recognition sequence. Conjugation was performed under Mn^2+^-dependent HUH reaction conditions at 37°C. Covalent DNA-protein conjugation was assessed by SDS-PAGE as a mobility shift relative to unconjugated HUH-aerolysin. EDTA was included as a negative control to chelate the divalent metal required for HUH catalysis. DNA containing multiple compatible HUH recognition sites generated multiple protein-DNA adducts.

For direct DNA labeling of cells, fluorescent oligonucleotide was conjugated to HUH-aerolysin, diluted to 500 nM, and incubated with U251 cells for 30 min. Cells were washed to remove unbound reagent and analyzed by fluorescence microscopy.

### RASUN cell-surface nuclease assay

The RApid Sensing of NUcleases (RASUN) probe consisted of a DNA hairpin containing an internal fluorescein fluorophore and BHQ-1 quencher:

TCTGTAAACTCGATGA/i6-FAMK/ATGGCCCGCAGCGACCACCCTTTGGGTGGTCGCTGCGGGCCATA/iBHQ-1dT/A

The hairpin was covalently conjugated to HUH-aerolysin using HUH-mediated DNA conjugation conditions. The assembled HUH-aerolysin–hairpin complex was diluted to a final concentration of 300 nM in complete cell-culture medium and added to cells following several PBS washes. Cells were incubated with the sensor for 60– 90 min and subsequently analyzed by flow cytometry.

For comparison of cell-surface nuclease activity, U251 and C2C12 cells were incubated with RASUN for 90 min. Sensor fluorescence was quantified by flow cytometry and compared with positive and negative sensor controls. C2C12 cells exhibited greater sensor activation than U251 cells under these conditions.

## QUANTIFICATION AND STATISTICAL ANALYSIS

Flow-cytometry data were analyzed using FlowJo. Median fluorescence intensity was used for quantitative comparisons unless otherwise indicated. Binding titrations were fit by nonlinear regression using **GraphPad Prism**. Data are presented as [**mean ± SD]**]. The number of independent biological replicates and statistical tests are indicated in the corresponding figure legends.

## Supplemental information

**Figure S1.**
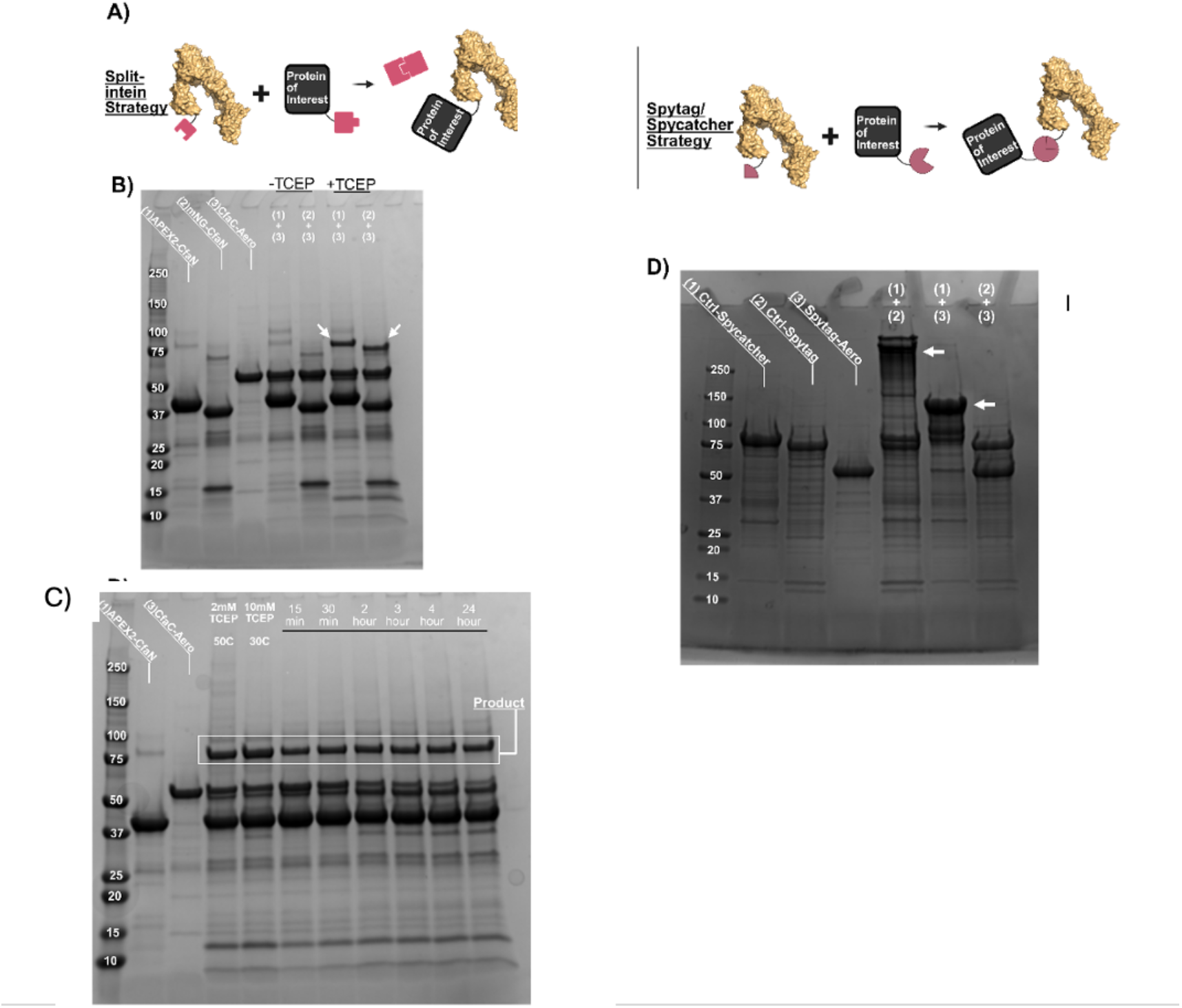
Evaluation of split-intein and SpyTag/SpyCatcher strategies for modular assembly of protein cargos with aerolysin. **(A)** Schematics of the split-intein and SpyTag/SpyCatcher approaches used to covalently couple independently expressed protein cargos to aerolysin. In the split-intein strategy, complementary intein fragments fused to aerolysin and a protein of interest associate and undergo protein trans-splicing to generate a covalently joined product. In the SpyTag/SpyCatcher strategy, complementary SpyTag and SpyCatcher modules spontaneously form a covalent linkage between aerolysin and the protein cargo. **(B)** SDS-PAGE analysis of split-intein-mediated ligation of the indicated aerolysin and cargo constructs in the absence or presence of TCEP. Arrows indicate products consistent with formation of the ligated species. **(C)** Optimization and time-course analysis of split-intein-mediated ligation under the indicated TCEP concentrations, temperatures, and reaction times. The boxed region indicates the product corresponding to the ligated protein complex. **(D)** SDS-PAGE analysis of SpyTag/SpyCatcher-mediated assembly of the indicated protein constructs. Arrows indicate higher-molecular-weight products formed upon incubation of complementary SpyTag- and SpyCatcher-containing proteins.

